# Detection of m6A from direct RNA sequencing using a Multiple Instance Learning framework

**DOI:** 10.1101/2021.09.20.461055

**Authors:** Christopher Hendra, Ploy N. Pratanwanich, Yuk Kei Wan, W.S. Sho Goh, Alexandre Thiery, Jonathan Göke

## Abstract

RNA modifications such as m6A methylation form an additional layer of complexity in the transcriptome. Nanopore direct RNA sequencing captures this information in the raw current signal for each RNA molecule, enabling the detection of RNA modifications using supervised machine learning. However, experimental approaches provide only site-level training data, whereas the modification status for each single RNA molecule is missing. Here we present m6Anet, a neural network-based method that leverages the Multiple Instance Learning framework to specifically handle missing read-level modification labels in site-level training data. m6Anet outperforms existing computational methods, shows similar accuracy as experimental approaches, and generalises to different cell lines with almost identical accuracy. We demonstrate that m6Anet captures the underlying read-level stoichiometry that can be used to approximate differences in modification rates. m6Anet achieves this without retraining model parameters, enabling the transcriptome-wide identification and quantification of m6A from a single run of direct RNA sequencing.

**Code Availability:** The source code for m6Anet is available at https://github.com/GoekeLab/m6anet. Installation instructions and online documentation is available at https://m6anet.readthedocs.io/en/latest/.

## Introduction

Modifications in RNA nucleotides were first discovered in the 1950s ^1,2^ in tRNAs and rRNAs, and today, more than 150 different modifications on RNA have been described ^3,4^. One of the most common RNA modifications is m6A which was discovered in 1974 as the main internal methylation on mammalian mRNA ^5,6^. This modification presents mostly at the consensus motif DRACH (D=A, G, or U, R=A or G while H is A, C or U) and has been shown to profoundly impact RNA structure ^7^, stability ^8,9^, splicing ^10^, and translation ^11^. Disruption of m6A homeostasis in animal models affects stem cell regulation ^12,13^, fertility and developmental process ^14^ while in humans, this modification plays an important role in cancer ^15,16^, cell-fate transition and determination ^17,18^ and transition, development ^19^, and diseases ^20,21^.

Experimental identification of RNA modifications can be achieved with three main approaches transcriptome-wide. Immunoprecipitation methods such as MeRIP-Seq ^22^, m6A-Seq ^23^, PA-m6A-Seq ^24^, m6A-CLIP/IP ^25^, miCLIP ^26^, m6A-LAIC-Seq ^27^, m6ACE-Seq ^28^, and m6A-Seq2 ^29^ use antibodies that specifically bind to the modified ribonucleotide. Chemical-based detection methods such as Pseudo-Seq ^30^, AlkAniline-Seq ^31^, utilise chemical compounds that selectively react with the modified ribonucleotide. Enzyme based approaches such as Mazter-Seq ^32^, m6A-REF-Seq ^33^ or DART-Seq ^34^ use specific enzymes to selectively distinguish modified and unmodified bases. The three approaches are similar in that they isolate the RNA after inducing changes to the surrounding nucleotides, followed by reverse transcription and sequencing using short read cDNA sequencing to detect these changes. While these approaches provide a transcriptome-wide map of RNA modification sites, they are limited by the availability of commercial antibodies and selective chemical reactivities for specific modifications ^35^, they lack single nucleotide resolution ^22,23^ and are incapable of identifying modifications for single RNA molecules.

The ability to sequence native RNA using Oxford Nanopore direct RNA-Seq can potentially overcome these limitations ^36^. Nanopore direct RNA-Seq infers the RNA sequence using the current intensity when an oligonucleotide passes through the pores. Modified nucleotides will emit a different signal intensity compared to unmodified nucleotides, allowing the computational identification of modified sites for each individual RNA molecule using either supervised or comparative approaches. Comparative approaches do not require training data for known RNA modifications but instead use control or reference samples to detect meaningful shifts in signal-based features that correlate to the presence of modifications. Comparative methods such as *Tombo* ^37^, *DRUMMER* ^38^, *nanoDOC* ^39^, *Nanocompore* ^40^, ELIGOS^41^, *xPore* ^42^, and *Yanocomp* ^43^ detect m6A sites by comparing with a sample with few or no m6A modifications. While these methods are accurate, their success relies on the availability of m6A-free control samples which typically involves silencing of specific writer genes which can be a limiting factor.

Supervised detection of m6A modifications involves training a classifier using labels that can either be obtained from synthetically modified RNA samples or existing experimental protocols such as miCLIP, MeRIP-Seq or m6ACE-Seq. Methods such as *EpiNano* ^44,45^, *MINES* ^46^, *nanom6A* ^47^, use training data to identify m6A using the sequencing error profile or shifts in the current signal intensity. Supervised methods can potentially be applied on a single sample, overcoming the main limitation of comparative methods for detection of specific RNA modifications. However, existing approaches are limited to a specific nucleotide content ^44–47^, and they are currently less accurate than comparative approaches using an m6A-free control ^40,42,43^.

One of the main challenges for supervised approaches applied to direct RNA-Seq data is that training data labels are provided for a set of reads at the site-level, but not for each individual read, which is known as a Multiple Instance Learning (MIL) problem in the machine learning literature ^48,49^. Existing methods address this problem by averaging read-based features ^44–46^. However, at any given site, we are likely to have a mixture of modified and unmodified reads and as such, not all reads provide useful features to detect m6A sites. Therefore, current approaches which do not consider the MIL structure in the training data might fail to detect m6A modifications from sites with low stoichiometry as it tends to obscure signals from the lowly expressed modified RNAs, and it limits the ability to integrate variation in read-level features into a predictive model.

To address these limitations we developed *m6Anet*, a MIL-based neural network model that takes in signal intensity and sequence features to identify potential m6A sites from direct RNA-Seq data. Our model takes into account the mixture of modified and unmodified RNAs and outputs the m6A-modification probability at any given site for all DRACH 5-mers represented in the training data. Unlike existing approaches, *m6Anet* learns high-dimensional representation of individual reads from each suspected site before aggregating them together to produce a more accurate prediction of m6A sites. By applying *m6Anet* to direct RNA-Seq data from different human cell lines we demonstrate that it is able to detect previously unlabelled m6A sites and also generalises across different cell lines without a reduction in performance. The approach utilized to train m6ANet is general enough that the network can be retrained to classify any natural or artificial RNA modifications given a set of labels.

## Results

### m6ANet identifies methylated positions with a multiple instance learning approach

Here we present *m6Anet*, a neural-network based Multiple Instance Learning model that combines learning the representation of each individual read with classifying m6a modified sites. *m6Anet* comprises two separate modules that are optimized jointly - a read level encoder and a pooling layer. The read level encoder uses signal and sequence features from each read, and transforms them into a high-dimensional representation before predicting the probability of each read being modified (Figure 1a). The read level probability is then pooled to give a probability estimate that a site is modified (Figure 1a). By combining features that represent signal and sequence properties, *m6Anet* can learn a model that can be applied for all 5-mers that are represented in the training data. Furthermore, the end-to-end training of our model implicitly learns a representation of the data that is optimized towards predicting the probability that a site is modified based on the assumption encoded within the pooling layer. In our case, the pooling layer represents the probability that a particular site contains at least one modified position, but in practice one can choose a pooling layer that best captures the labelling process associated with the data collection step. While we apply *m6Anet* to the task of m6A RNA modification detection, the framework generalises to any other task for which training labels are available, such as DNA modification detection or other RNA modifications of interest. *m6Anet* is implemented in Python and available through GitHub (https://github.com/GoekeLab/m6anet).

**Figure 1.**
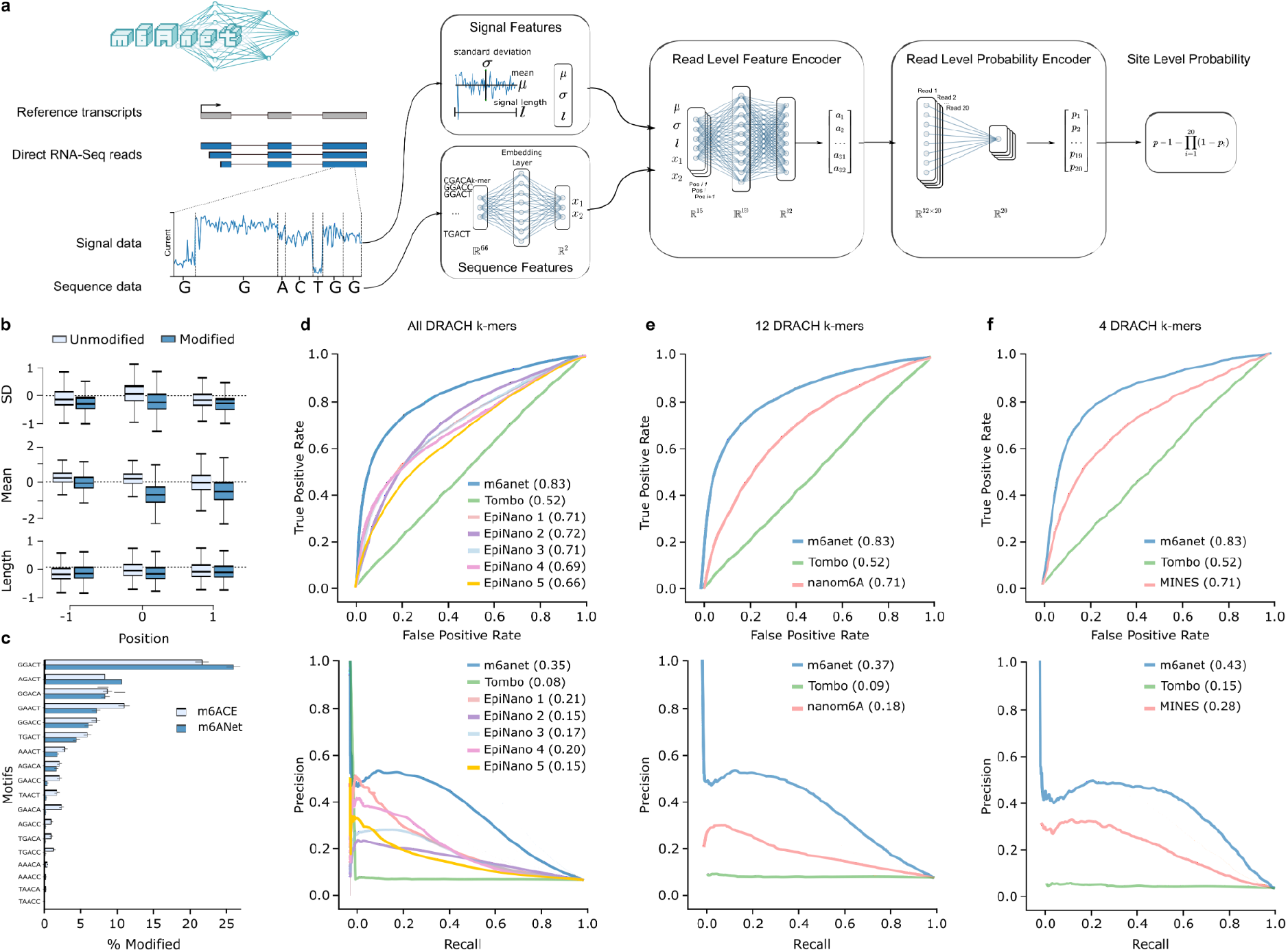
Schematic of m6Anet and evaluation on detection of m6A in human cell lines. (a-b) m6Anet model schematics. (b) Box plot showing the difference in average features distribution between different *m6Anet* prediction. The horizontal lines on the boxes show median, Q1, and Q3 and 1.5 interquartile range (c) Comparison of the proportion of modified sites predicted as modified by *m6Anet* and by m6ACE on the top 4 modified 5-mers (GGACT, GAACT, GGACA, AGACT). (d) ROC Curve and PR Curve of *m6anet* against all 5 EpiNano models and Tombo. (e) ROC Curve and PR Curve of *m6anet* against *nanom6A* and *Tombo*. (f) ROC Curve

### Training data for m6Anet model parameter estimation

To learn the model parameters, *m6Anet* requires training data consisting of labels (modified/unmodified) and direct RNA-Seq reads. In order to train a model for m6A we used labels obtained from m6ACE-Seq that identifies m6A at single nucleotide resolution ^28^. *m6Anet* uses positions which are identified to have m6A as labels for the *modified* class, and any other position with the same 5-mer sequences that are included in the modified class will be used as the *unmodified* class. Since m6A modifications occur at the DRACH motifs, we removed any non DRACH motifs from these data for *m6Anet*, however, this step is not required for training data without prior knowledge about the motifs. Since m6A modifications are rare compared to unmodified sites, we oversample the modified sites during training to obtain a balanced data set. Here, we used direct RNA-Seq data from the HCT116 cell line for which matched m6ACE-Seq data is available as part of the Singapore Nanopore Expression Project ^50^.

### Contribution of signal and sequence features to m6Anet predictions

*m6Anet* uses signal features corresponding to the normalised signal intensity, standard deviation, and dwelling time for each position. To understand how each feature contributes to the prediction of *m6Anet*, we explored the difference in features distributions between the predicted modified sites and predicted unmodified sites for each one of the DRACH motifs. Signal intensity of the center base pair showed the strongest difference between predicted modified and predicted unmodified sites, with dwell time showing the smallest difference in distributions (Figure 1b, Suppl. Figure 1a). However, all features distinguish modified and unmodified sites and are informative for m6A predictions.

As RNA modifications can affect the nanopore current signal at the neighbouring bases, we tested whether information from additional positions increases the model accuracy. We performed 5 fold cross validation with features extracted from 0 to 5 base pairs flanking the candidate sites to evaluate the additional value of neighbouring positions, splitting the data at the gene level to ensure independence between training set and test set. Our results show that *m6Anet* performance is highest when 1 base pair flanking positions were considered, whereas additional information from the neighbouring features beyond 1 base pair did not result in any further improvement of the classifier (Supplementary Table 1).

A key feature of *m6Anet* is the ability to jointly model RNA modifications for all candidate 5-mer sequences in the training data. To evaluate if this approach biases the prediction of m6A sites based on the sequence, we compared the 5-mer frequency of predicted m6A sites with the 5-mer frequency observed in m6ACE-Seq data on positions that have not been used to train *m6Anet* model parameters. We find that *m6Anet* predictions have a comparable 5-mer profile as the m6ACE-Seq data, with less frequent motifs being equally represented (Figure 1c, Suppl. Figure 1b), showing that *m6Anet* captures the expected modification rates per 5-mer from a single model that combines features from signal and sequence.

### m6Anet accurately identifies m6a sites from direct RNA-Seq data

To evaluate the performance of *m6Anet* we tested the model on direct RNA-Seq data from the HEK293T cell line ^42^, using m6ACE-Seq ^28^ and miCLIP data ^26^ from the same cell line as ground truth. Using these data, we compared the performance of *m6Anet* against *EpiNano* ^44,45^, *MINES* ^46^, *Tombo* ^37^, and *nanom6A* ^47^ using the Area Under the Curve (AUC) of the Receiver Operating Characteristic (ROC) and Precision Recall (PR) curves to quantify the model accuracy. On the HEK293T cell line, m6Anet achieves a ROC AUC of 0.83 and PR AUC of 0.35 (Figure 1d, Supplementary Table 2). Among the other methods, only *EpiNano* and *Tombo* return predictions for all DRACH motifs, however, at a lower accuracy compared to *m6Anet* (*EpiNano*: ROC AUC: 0.69-0.72, PR AUC: 0.15-0.21; *Tombo*: ROC AUC:0.52, PR AUC: 0.08) (Figure 1d). Since *MINES* and *nanom6A* output predictions only for 4 and 12 5-mers respectively, we ran separate validation between *MINES*, *nanom6A*, and *m6Anet* on these motifs alone. On these data, *m6Anet* achieved a ROC AUC of 0.83 (4 motifs and 12 motifs) and a PR AUC of 0.43 (4 motifs) and 0.37 (12 motifs) outperforming both *MINES* (ROC AUC: 0.71; PR AUC: 0.28) and *nanom6A* (ROC AUC:0.71; PR AUC:0.18) (Figure 1e,f, Supplementary Tables 2), suggesting that *m6Anet* provides the most accurate predictions of candidate m6A among existing methods.

### Novel m6Anet predictions are sensitive to METTL3 knockout

While the overall accuracy for detection of m6A from direct RNA-Seq data is high, many m6A sites predicted by *m6Anet* are not identified by these experimental approaches. Different methods for profiling m6A have been described to identify different sets of m6A sites ^28^. Indeed, in the HEK293T cell line, the largest number of sites are detected by only one protocol (Figure 2a). Among the three protocols, *m6Anet* predictions show an equal or higher fraction of support by other technologies, suggesting that *m6Anet* is comparable to existing experimental protocols (Figure 2b).

**Figure 2.**
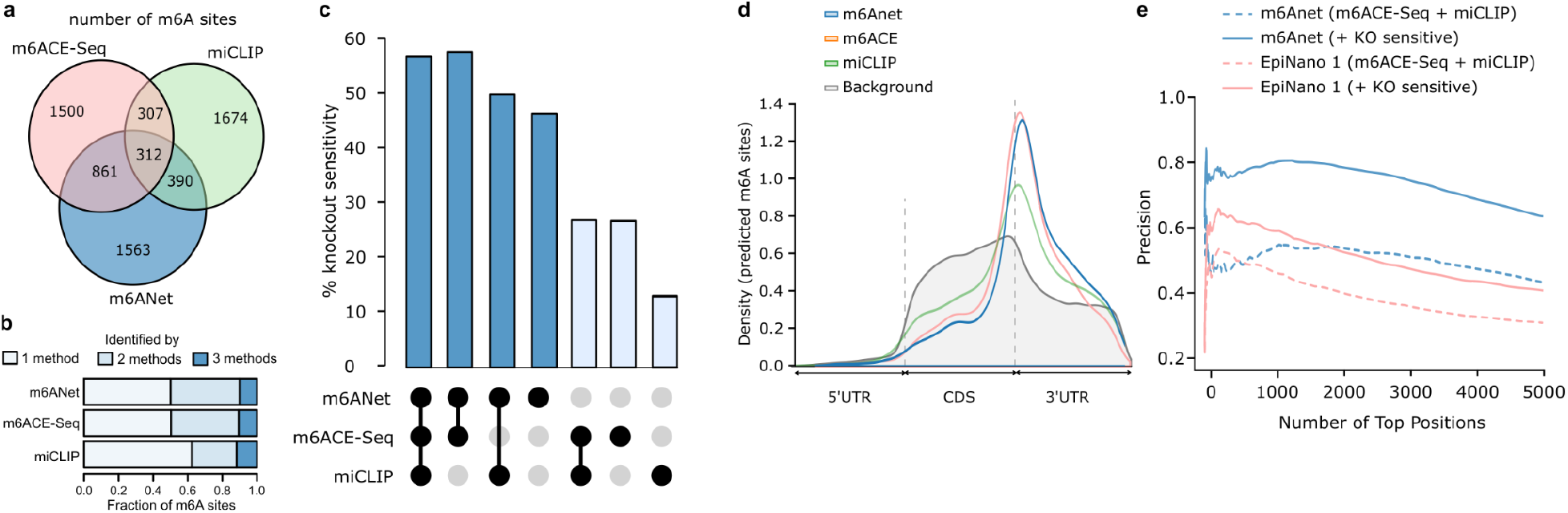
Performance comparison between *m6Anet*, m6ACE-seq and miCLIP on HEK293T cell line. Experimental design: Comparison is done with labels obtained from m6ACE and miCLIP on all DRACH positions (a-b) Total number of modified sites captured by *m6Anet*, m6ACE-seq and miCLIP (c) Percentage of captured sites that show significant shift in signal distribution against METTL3-KO for each of the three protocols (d) Metagene plot of the modified sites captured by the three protocols against the background distribution of all DRACH sites in the data that has at least 20 reads (e) The adjusted true positive rate after including position sensitive to METTL3-KO of *m6Anet* and *EpiNano*.

In order to evaluate whether the novel sites predicted by *m6Anet* are valid m6A sites, we identified positions which are sensitive to loss of the m6A writer METTL3. Using an existing comparative approach (xPore), we mapped m6A sites in the HEK293T cell line by comparing it against a METTL3 knockout cell line that is depleted of m6A ^28,42^. We then define DRACH sites which have a significant difference compared to this control as knockout sensitive sites (KO sensitive), resulting in 1888 candidate positions when a stringent threshold is used (Suppl. Table 2, see methods). The sites which are detected by all three methods show the highest fraction of KO sensitive sites (57% Figure 2c). Among the sites which are only detected by one method, *m6Anet* predictions have the highest proportion of KO sensitivity detected by *xPore* (46%, Figure 2c), with a less stringent method to define KO sensitive sites further increasing the fraction for all 3 protocols (Suppl Figure 2a). As the usage of a direct RNA-Seq based method for evaluation might favour *m6Anet* predictions, we also investigated the enrichment of m6A positions along the transcript coordinates. This analysis shows that all the sites that are captured by the three methods are enriched in the 3’ end of the CDS as expected for m6A (Figure 2d). m6A sites which are only found in one method show a similar pattern, with m6ACE-Seq and *m6Anet* predictions showing the strongest enrichment (Suppl Figure 2b), suggesting that many of these are indeed valid m6A positions that are only detected by a single technology.

Including the additional METTL3 KO sensitive m6A sites into the validation set increases the estimated precision for *m6Anet* and other methods based on direct RNA-Sequencing (Figure 2e, Suppl. Figure 2c, Suppl. Table 2). These results confirm that many novel m6A sites identified by m6Anet are sensitive to METTL3 loss and that the true precision of *m6Anet* is underestimated when comparing it to labels obtained from miCLIP or m6ACE-Seq, most likely reflecting technology-specific m6A predictions.

### m6Anet generalises to new cell lines without loss in accuracy due to training

In order to test how well *m6Anet* generalises to data from a new cell line, we compared the models trained on the HCT116 and HEK293T cells. For this comparison, we split the dataset on the gene level into a training and test set, ensuring that the test sets on both cell lines comprised the same genes (Figure 3a, b). We find that both models trained on reads from HCT116 and HEK293T respectively generate predictions with a similar accuracy when applied on the same cell line (Figures 3c,d, Suppl. Figures 2d-e, Supplementary Tables 3,4). On the HCT116 cell line, the model learned on the HEK293T data even shows a better performance than on the original cell line used for training (Figure 3c,d). Furthermore, both models were able to identify m6A sites on genes which are not expressed in the cell lines they were trained on (Figure 3e,f) demonstrating that *m6Anet* generalises to other cell lines without a loss in accuracy due to cell type-specific training data.

**Figure 3.**
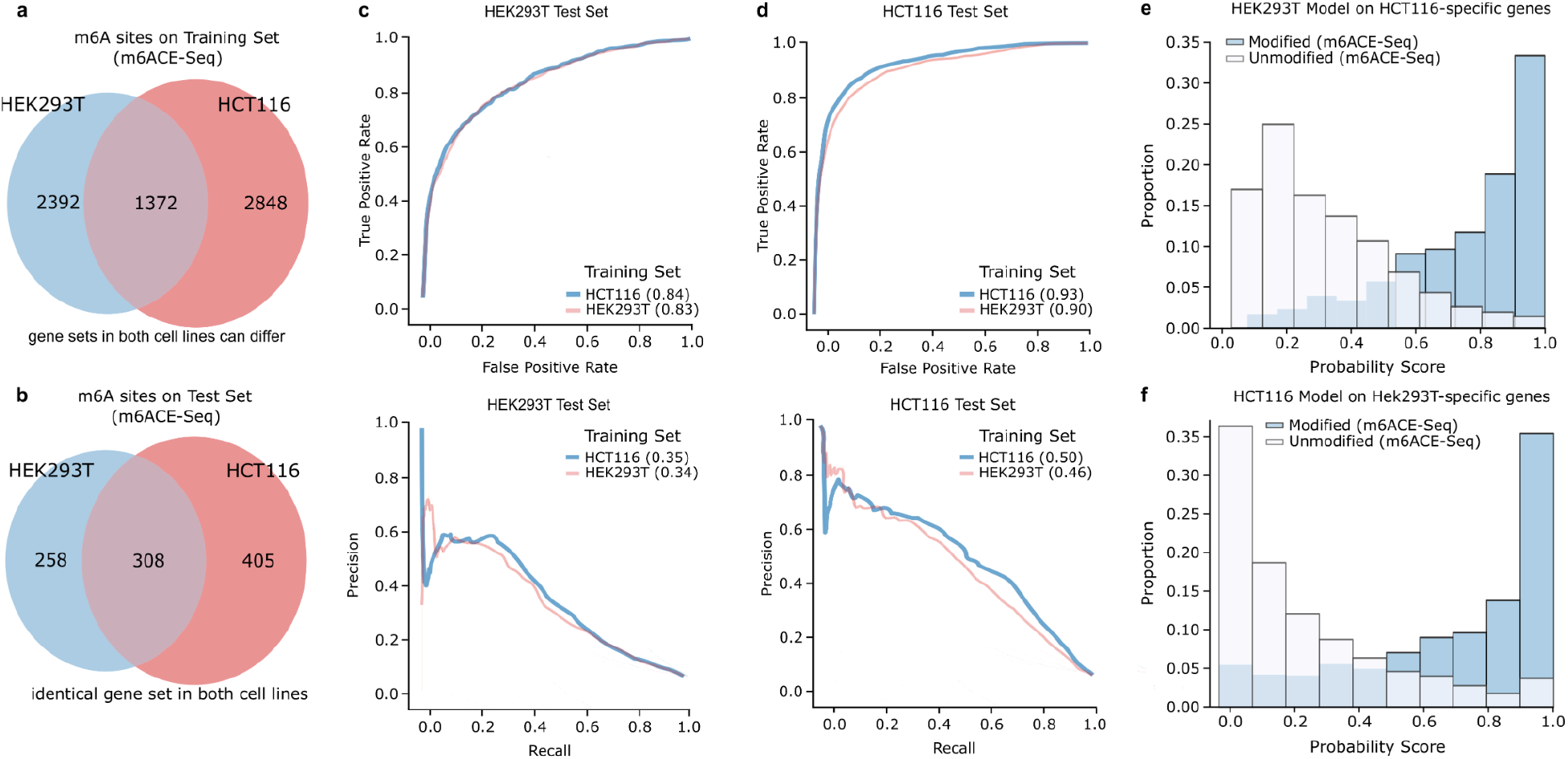
Comparison of *m6Anet* model across two different cell lines. Experimental design: Comparison is done with labels obtained from m6ACE on HCT116 cell line and both m6ACE and miCLIP for HEK293T cell line. We split each cell line into training set and test set on the gene level with the test set selected from common genes across the two cell lines to ensure that each model is not trained on a set of genes used for training.(a-b) Distribution of modified positions across both cell lines on the training sets and the test sets. (c) ROC Curve and PR Curve of the models trained on the HCT116 train set and HEK293T train set on the HEK293T Test set (d) ROC Curve and PR Curve of the models trained on the HCT116 train set and HEK293T train set on the HCT116T Test set (e-f) Distribution of probability score of HEK293T (HCT116) model on the genes that are expressed only on the HCT116 cell test set (HEK293T test set). Histogram shows that *m6Anet* trained on both cell lines can make accurate predictions on a set of genes that are not present in their original training data

### m6Anet provides single molecule m6A predictions

While the primary output of *m6Anet* is a site-level modification probability, it was designed to learn a hyper-dimensional representation of each read based on its signal and sequence features, which is then used to infer a read-level modification probability. This design allows the identification and visualisation of modifications for individual RNA molecules at candidate m6A sites. To illustrate the ability to predict per molecule modification status, we extracted the read level representation and probabilities from both the HEK293T wild type and knockout cell lines for candidate m6A positions (p>0.9 in wild type cells, p<0.2 in knockout cells, see methods). We then performed a Principal Component Analysis (PCA) on the read-level features to map reads into a 2-dimensional space. We find that reads form two clusters that are dominated by the knockout reads (unmodified cluster) and wild type reads (modified cluster) (Figure 4a, Suppl. Figure 3a-k). Using these clusters we projected data from individual reads for the positions identified to have the highest modification probability into this read-level feature map (Figure 4b, Supplementary Table 5). While reads from the knockout sample have low predicted m6A probabilities and fall into the knockout cluster, reads from the wild type samples are enriched in the cluster with high m6A probability, providing insights into the single molecule predictions by *m6Anet* (Figure 4c, Suppl. Figures 3l-n).

**Figure 4.**
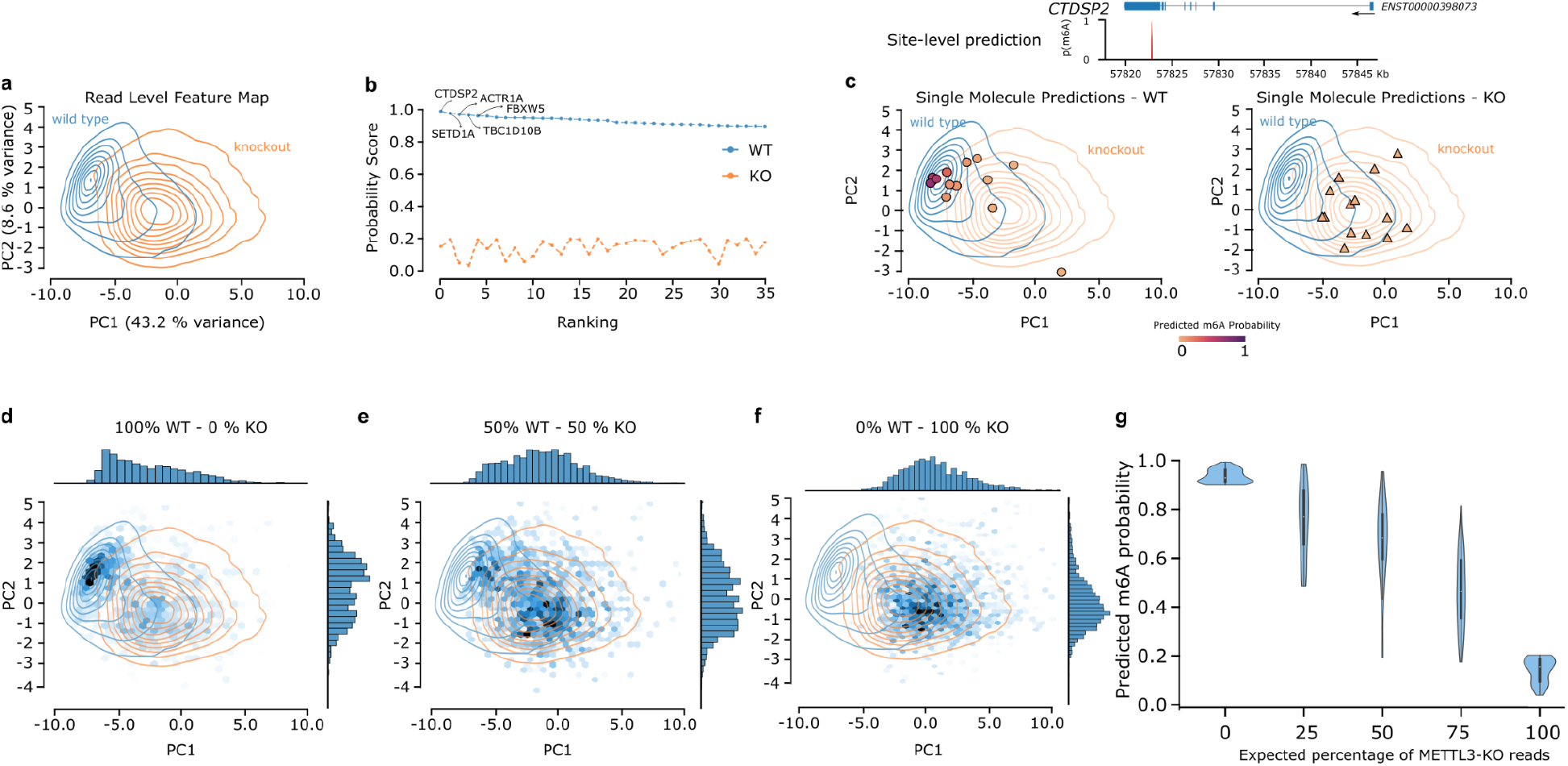
Quantification of m6Anet on HEK293T cell line. Experimental design: We sample 100 reads from each DRACH sites and extract the output from the second last layer of *m6Anet* for each of these reads and visualize them on two dimensional space using PCA (a) Density plot of the top 4 modified 5-mer (GGACT, GAACT, GGACA, AGACT) for both Wild Type and Knockout sample after filtering for positions that are almost 100% modified on the Wild Type sample (p >= 0.9) and almost 0% modified on the KO sample (p <= 0.2) (b) Ranking plot of the positions in (a) and the genes associated with the top positions (c) Scatter plot of 20 randomly sampled reads from the top ranked position(d-f) Hex plots of the read level feature map for 0%, 50%, and 100% KO mixtures on filtered positions. Changes in the concentration of points as visualized on the first two principal components of the same PCA space as in Figure 4. The gradual shifts from (d) to (f) suggests that m6Anet read features capture the expected change in the stoichiometry of m6A modifications. (e) Violin plot of the probability score of the top predicted positions by m6Anet across the 5 mixtures. The plot shows an expected decrease in the predicted m6A probability as the percentage of METTL3-KO reads increases.

### Site-level m6A probabilities capture differences in m6A stoichiometry

As *m6Anet* integrates read level probabilities to obtain the final site level probability, these observations suggest that it might reflect the underlying modification stoichiometry. To validate whether a change in the proportion of modified reads is reflected in a change in site-level m6A probabilities, we analysed direct RNA-Seq data from METTL3 knockout and wild type samples that were mixed at specific proportions corresponding to an expected relative m6A stoichiometry of 0%, 25%, 50%, 75%, and 100% ^42^. On the set of sites which were predicted to be modified in the 100% wild type samples (p>=0.9) and which are predicted to be unmodified in the knockout samples (p<=0.2), we observed a gradual shift of reads from the modified cluster to the unmodified cluster, corresponding to the expected changes in the relative m6A stoichiometry (Figure 4d-f, Suppl. Figures 4a-b). Similarly, the m6A site-level probability predictions are reduced corresponding to a reduction in expected modification rates on the same set of sites (Figure 4g; Suppl. Figures 4c,d, Supplementary Table 6), further suggesting that these probabilities reflect the change in the proportion of modified reads. While the primary purpose of the site-level probability is to provide an estimate of confidence, these data suggest that it captures variation in the underlying modification rates that can be used to compare sites within one sample, or to estimate global differences in m6A abundance across multiple samples or conditions.

## Discussion

Supervised approaches promise to enable the accurate detection of RNA modifications from direct RNA-Seq data. These methods rely on accurate training data, which can be obtained through experimental protocols such as m6ACE-Seq or miCLIP, or through synthetic data. However, experimental methods only provide site-level modification labels, whereas Nanopore data is provided for individual RNA molecules for which the modification status is not observed. Here we address this by developing *m6Anet*, a neural-network based Multiple Instance Learning model. *m6Anet* combines learning the representation of each individual read with classifying m6a modification sites, outperforming other existing computational methods and providing an accuracy that is comparable to experimental approaches.

Even though *m6Anet* was designed to handle missing read-level modification information, it still relies on the accuracy of site-level modification training data. Depending on how these data were generated, such labels could be incomplete ^51,52^, or include multiple distinct modifications ^26,28^ thereby introducing noise in the training data and a reduction in the model performance. Here we find that the prediction accuracy on m6A appears to be high even when different training data sets are used. Nevertheless, additional training data on different modifications, species, and experimental protocols will likely further improve the prediction accuracy for supervised approaches such as *m6Anet*.

While supervised methods can identify RNA modifications in a single sample, comparative methods facilitate the analysis across conditions ^40,42,53^. However, one of the key advantages of supervised methods over comparative methods is their ability to predict the occurrence of specific RNA modifications such as m6A. By predicting m6A modifications on candidate sites identified by comparative methods, *m6Anet* can overcome their inability to assign specific modification types, thereby facilitating modification-specific analysis of differential modifications.

In contrast to short-read based experimental approaches for profiling RNA modifications, direct RNA-Seq is a simple assay that can make m6A profiling scalable. However, similar to experimental protocols which are influenced by aspects such as antibody-specificity ^26,28^ the accuracy of m6Anet will be influenced by aspects such as the sequencing chemistry, basecalling algorithms or accuracy in the alignment of reference sequence to signal. Improvements in the sequencing technology and methods that extract summarised data from Nanopore signals can further increase the accuracy of *m6Anet*. While we observe a high number of technology-specific m6A predictions, our data supports that these are likely valid m6A sites, suggesting that *m6Anet* and short read-based methods already have a comparable accuracy in detection of m6A.

Here we applied *m6Anet* to identify m6A modifications, however it was designed to facilitate training on any RNA modification phenotype of interest. While *m6Anet* could be used to identify other naturally occurring RNA modifications, it can also be trained to predict artificial modifications that help to identify single molecule RNA structures ^54^. A key advantage of direct RNA-Sequencing is the ability to profile the modification status of individual reads. While the evaluation of single molecule predictions is still limited due to the inability to generate single molecule reference data, our analysis suggests that *m6Anet* single molecule predictions correspond to the expected global modification rate. As *m6Anet* generalises well to new data, it can be directly used for the standalone identification of m6A and possibly other modifications after retraining. However, it will also complement existing experimental approaches by increasing confidence and resolution, enabling the accurate site level modification prediction while facilitating the additional exploration of single molecule modification probabilities from a single run of direct RNA-Seq data.

## Methods

### *m6Anet*: a Multiple Instance Learning based Neural Network

*m6Anet* performs RNA modification detection using direct RNA-seq data by formulating it as a Multiple Instance Learning problem. Each position *i* corresponds to a k-mer sequence *S* of length *k* = 5 with:

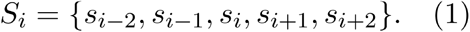

Here *s_i_* ∈ {A,C,G,U} corresponds to the nucleotide of position *i*. For each position, the site modification status is given by *y_i_* where

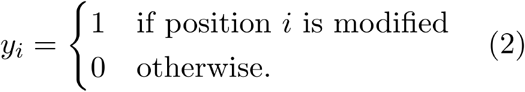

We also assume that each read *j* at position *i* has a modification status described by *y_i,j_* given by:

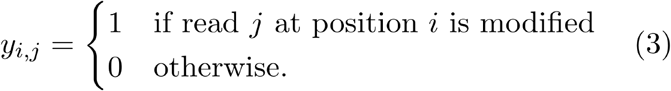

While *y_i_* can be observed, *y_i,j_* cannot be observed and remains unknown. Each read *j* at position *i* is described by the feature vector *x_i,j_* ∈ ℝ^15^ with:

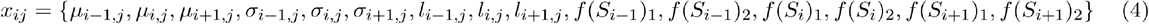

where *μ_i,j_* represents the normalized mean nanopore raw signal of read *j* at position *i*, *σ_i,j_* represents the normalized standard deviation of the nanopore raw signal of read *j* at position *i*, and *l_i,j_* represents the normalized dwelling time of read *j* at position *i*. Furthermore, we encode all *N_S_* possible 5-mer sequence motifs *S* that are included in the training data into a 2-dimensional vector using a neural network embedding layer *f* : *N_S_* → ℝ^2^, with *N_S_* = 66 in the case of m6A (DRACH). Thus, the quantity *f*(*S_i_*)_*k*_ gives the k-th dimension of the embedded vector of the 5-mer motif *S_i_*, with *k* ∈ {1, 2}. Each position *i* with *N_i_* reads is then described by

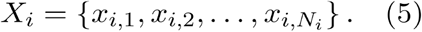

In the first step, m6Anet estimates the *read level modification probability p_ij_* of the read *j* at position *i* being modified:

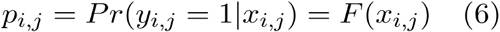

where *F* : ℝ^15^ → ℝ is parameterized by a neural network with two hidden layers of dimension 150 and 64 respectively. In the second step, m6Anet pools the read level probability using a noisy-OR pooling layer to estimate the *site level modification probability P_i_*:

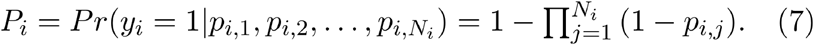

The noisy-OR pooling layer captures the assumption that a site is modified if at least one of its reads is modified. In practice, the noisy-OR pooling layer encourages any gradient-based learning methods to update the model parameters with respect to all reads instead of just a single modified reads. As a result, the site probability estimated by m6Anet should reflect the changes in the number of modified reads between different sites.

To train the network, we minimize the average cross entropy loss 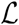 between *P_i_* and *y_i_* for all sites

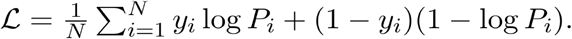

Here *f* and *F* are learnt in an end-to-end fashion by minimizing the cross entropy loss *L* with the Adam optimizer. Consequently, the network learns to predict the individual read probability *p_i,j_* along with optimized sequence representation *f*(*N^S^*) that will minimize the discrepancy between *P_i_* and with *y_i_* respect to the noisy-OR pooling layer. We have evaluated alternative pooling layers, such as the Attention and gated Attention-based pooling ^55^ but have not found any statistically significant improvement in the performance of *m6Anet* compared to the noisy-OR pooling layer for m6A detection.

### Preprocessing for m6Anet

m6Anet requires the output from *Nanopolish eventalign* function ^56^ in order to group continuous Nanopore current measurements from each read into events and map them to their corresponding positions in the transcriptome. Each nanopolish event comprises the mean, standard deviation, and dwelling time of its constituting raw signals and since multiple events can be assigned to the same location in the transcriptome, *m6Anet* then takes a weighted average of each of these features based on the size of their respective groups. Afterwards, *m6Anet* discards positions with mismatched 5-mers and computes the mean and standard deviation of the signal features for each possible 5-mer motif across the transcriptome. Lastly, *m6Anet* performs z-normalization on the weighted average features based on the mean and standard deviation of the 5-mers motif of the given segment. The preprocessing function is implemented in *m6Anet*.

### Data Processing

#### Processing of direct RNA sequencing data

All data used in this work was obtained from ^42^, ^50^. To train and validate *m6Anet*, we downloaded a single replicate (replicate 2 run 1) of the HCT116 cell line and a single replicate of the HEK293T cell line (replicate 1) while to run *xPore*, we downloaded all replicates of the HEK293T cell lines as recommended. Data was basecalled from the raw fast5 files using Guppy and aligned to the transcriptome with minimap2.1 (minimap2 ‘-ax map-ont -uf–secondary=no’) using the GRCh38 Ensembl annotations release version 91. We used a combined FASTA file containing coding and noncoding RNA reference annotations, keeping only the transcripts that matched the reference genome annotations (nf-core/nanoseq:https://doi.org/10.5281/zenodo.3697960). Afterwards, we ran *Nanopolish 0.11.3* with the --scale-events and --signal-index options.

#### m6A-cross-linking-exonuclease sequencing

Modified positions for m6ACE-seq are obtained from ^28^, ^42^ where we also follow their preprocessing steps for the HEK293T cell lines and include only those positions that are METTL3-dependent (WT/KO relative methylation level ratio ≥4.0, *P* value of one-tailed *t*-test, <0.05). As for the HCT116 cell line, we consider any sites that appear in the m6ACE-seq library to be modified since the absence of METTL3-KO data means we are not able to filter based on the WT/KO relative methylation level like in the HEK293T cell lines.

#### m6A individual-nucleotide-resolution cross-linking and immunoprecipitation

Modified positions from miCLIP were obtained from ^26^ where we combine both CIMS and CITS miCLIP libraries from the supplementary and consider a position to be modified if it is found in any of these libraries.

### Model Evaluation

#### Contribution of Flanking Regions to m6Anet Performance

In order to evaluate the performance of *m6Anet* under different combinations of features, we performed a 5-fold cross validation on the HCT116 dataset. In each fold, we train our model on 75% of our training data for 60 epochs and choose the model that performs the best on the remaining 25% of the training data and validate the performance of the model on the test set. We also ensure that no genes are shared between the train, validation, and test set during the evaluation. During training the parameters of the model are learnt by minimizing the cross entropy loss using the Adam optimizer ^57^ with amsgrad ^58^ turned on. On each site, we sample 20 reads and during test time, we run the model 5 times and average the probability value across the 5 runs. Results are shown in Supplementary Table 1. All models are implemented on Pytorch v1.7.1 ^59^. Training is done with a fixed learning rate of 0.0004 and a mini-batch size of 512 on a single NVIDIA GeForce GTX 1080 Ti.

#### Comparison between m6ANet and other models on HEK293T Cell Line

In order to have a fair comparison between *m6Anet* and existing methods to detect m6A modifications, we performed the comparison against other models on the HEK293T cell line which was not used to train the *m6Anet* model. We consider a position to be modified if it is captured by either miCLIP or m6ACE-Seq as modified and we only consider DRACH sites that have at least 20 reads.

##### Tombo

We ran *Tombo version 1.5.1* from https://github.com/nanoporetech/tombo. To detect modifications, we first resquiggled the raw reads with *tombo-resquiggle* and performed de-novo detection with *tombo detect_modifications de_novo*. Since *tombo* outputs a fraction of modified reads per position, we treat this as the probability of a site being modified for our comparison.

##### EpiNano

We ran *EpiNano 1.1 and 1.2* from https://github.com/enovoa/EpiNano and in both cases, we excluded feature generations for positions that do not contain AC center nucleotides (without this step, the results were not returned within 7 days on a AMD EPYC 7R32 server with 180GB of memory). There are 4 SVM models on *EpiNano 1.1* and 1 SVM model on *EpiNano* 1.2 that could work with a single sample of direct RNA sequencing data. We numbered these models from 1 to 5 respectively.

##### MINES

We ran *MINES* from https://github.com/YeoLab/MINES on cDNA mode, following the steps that are specified in the readme file on the github page. The original *MINES* model does not output the probability of a site being modified but instead only shows sites that are considered modified. For this comparison, we modified the code so that the RandomForest model outputs the probability of a site being modified and we compared the results with m6Anet on sites shared between the two methods. The modified code is available at https://github.com/chrishendra93/MINES.git.

##### nanom6A

We ran nanom6A from https://github.com/gaoyubang/nanom6A. Similar to *Tombo*, it only outputs a fraction of reads that are modified for each site and so we treat these numbers as the probability of a site being modified.

### Comparison between m6Anet, m6ACE-Seq, and miCLIP

In order to evaluate the relative performance between m6Anet and other commonly used experimental protocols, we performed a comparison with miCLIP and m6ACE on the HEK293T cell line. We set a P=0.9 threshold for m6Anet site probability to select modified sites. miCLIP and m6ACE-Seq data was obtained and processed as described above.

To calculate whether a site is knockout sensitive or not, we ran *xPore 1.0* on replicate 1, 2, and 3 of the HEK293T samples provided by ^42^ with pooling option and a minimum read threshold of 20. To be conservative about our estimates, we imputed any sites that are not present in the *xPore* run with *P* value of 1 (not differentially modified). We performed multiple test corrections using Benjamini-Hochberg procedure and set an alpha rate of 0.05.

To obtain a second (less stringent and less accurate) estimate for knockout sensitive sites we also ran Welch’s t-test from the scipy package’s function ttest_ind (setting equal variance to false). Similar to the analysis with *xPore*, we pooled reads from all three replicates and required tested positions to have a minimum of 20 reads. We then performed multiple test corrections using Benjamini-Hochberg procedure, set an alpha rate of 0.05 and imputed any other sites that do not meet the filter criteria with a P value of 1.

#### Metagene plot

To visualise the distribution of m6A sited across the transcript (metagene plot), we first mapped each gene coordinate to transcript coordinate based on the most expressed transcripts per gene. Afterwards, we annotate each position based on its location along the transcript as 3’UTR, 5’UTR, or coding sequence. We then calculate the relative position of each position on the transcript and plot the abundance of those positions that are considered modified by *m6Anet*, m6ACE-seq or miCLIP.

### Comparison of m6Anet performance on HEK293T and HCT116 cell lines

In order to measure the robustness of *m6Anet* across different cell lines, we train two different models on the HEK293T and HCT116 cell lines respectively and measure the performance of each model on both HEK293T and HCT116 test sets. We randomly select 500 genes that are present in both cell lines to form two test sets for both cell lines and use the remaining genes as training data. We further split 20% of the training set for each cell line at the gene level into a validation set for model selection.

### Visualisation of single molecule modification probabilities

#### Principal Component Analysis and Read Level Feature Map

In order to learn the read level feature map that visualises single molecule m6A probability predictions, we project the high-dimensional read representations of *m6Anet* using a Principal Component Analysis and visualize the first two principal components. We sampled 100 reads from each position and extracted the 64-dimensional features generated by the second last layer of *m6Anet* from each of these reads. We ran PCA from the python package *scikit-learn* ^60^ with n_components set to 0.99 and svd_solver set to full so that the algorithm will choose the number of components that will result in total variance explained to be as close as possible to 1.

To better visualize the features that are representative of both modified and unmodified reads, we first filtered for positions that are highly modified in the WT sample (P >= 0.9) or unmodified in the KO sample (P <=0.2) and which contain the 5-mer motifs GGACT, GAACT, GGACA, or AGACT. These motifs are chosen because they represent the most modified 5-mer motifs in the HEK293T cell lines based on miCLIP annotations or m6ACE-seq annotations. We further sampled 20 reads from each of these positions in order to minimize running time. We then calculated the density plot and hex plot on both the wild type reads and knockout reads on the first two principal components of the read features using Python seaborn package. We then use the resulting density plot as a read level feature map to visualise individual molecule modification probabilities.

### Quantification of m6Anet on HEK293T mixtures

#### Analysis of wild-type - METTL3 knockout mixture samples

To analyse the ability of *m6Anet* to estimate m6A stoichiometry we used the Wild Type - METTL3 knockout mixtures from ^42^ that have an expected relative average modification rate of 0% (METTL3 knockout), 25%, 50%, 75%, and 100% (wild type). We filter for those positions that are present in all samples and are either fully modified (probability greater than 0.9 in the 100% Wild Type sample) or not modified (probability less than 0.2 in the KO samples).

### Data Availability

The HCT116 cell lines data were obtained from the Singapore Nanopore Expression Project ^50^ through ENA (PRJEB44348) while the HEK293T cell lines data along with its KO variants and KO mixture variants were obtained from ^42^ through ENA (PRJEB40872).

## Acknowledgements

C.H. is supported by funding from the Institute of Data Science, National University of Singapore and NUS Graduate School - Integrative Sciences and Engineering Programme (ISEP). J.G. is supported by funding from the Agency for Science, Technology and Research (A∗STAR), Singapore, and by the Singapore Ministry of Health’s National Medical Research Council under its Individual Research Grant funding scheme.

## Competing Interests

J.G. received reimbursement for travel and accommodation from Oxford Nanopore Technologies to present at the Nanopore Community Meeting in San Francisco in 2018.

## Supplementary Tables

- **Supplementary Table 1.** Cross validation results on the HCT116 cell line with 0 to 5 base pairs neighboring positions on 3 different model selection criteria (average loss, best ROC AUC and best PR AUC). Columns show the accuracy as measured by the Area under the ROC Curve (roc_auc) and the PR Curve (pr_auc).
- **Supplementary Table 2.** Predicted modification probabilities on the HEK293T cell line on the 18 DRACH motifs by m6Anet, Tombo, EpiNano, MINES, and nanom6A. Columns show the individual probability score by each model along with the adjusted *P* value given by xPore and t-test, labels from m6ACE-Seq and miCLIP.
- **Supplementary Table 3.** Probability scores of models trained on HCT116 cell line and HEK293T cell line on the HCT116 test set. Columns show the individual probability score of each model and the modification status value of 1 indicates that the site is modified while 0 indicates that the site is not modified based on m6ACE-Seq data.
- **Supplementary Table 4.** Probability scores of models trained on HCT116 cell line and HEK293T cell line on the HEK293T test set. Columns show the individual probability score of each model and the modification status value of 1 indicates that the site is modified while 0 indicates that the site is not modified based on m6ACE-Seq data.
- **Supplementary Table 5** Probability scores of sites shared by the wild type and knock out variants of the HEK293T cell lines. Columns show the transcriptomic and genomic coordinates along with the probability scores of each sample and the 5-mer motifs of each position
- **Supplementary Table 6** Probability scores of sites shared by the wild type and mixtures of knock out variants of the HEK293T cell lines. Columns show the transcriptomic and genomic coordinates along with the probability scores of each sample and the 5-mer motifs of each position

---

**Supplementary Figure 1.**
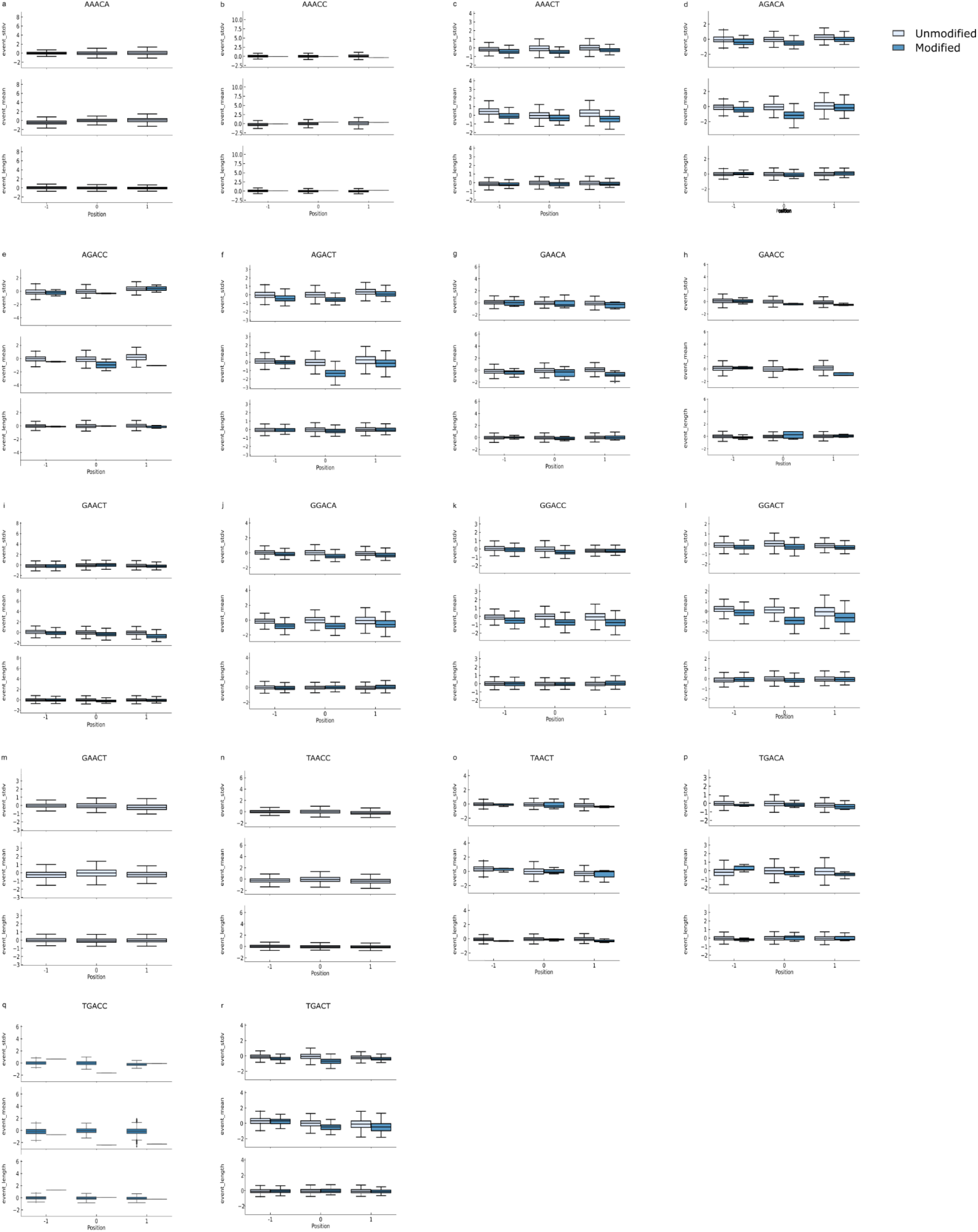
(a) Box plot showing the difference in average features distribution between different *m6Anet* prediction across all 5-mers. The horizontal lines on the boxes show median, Q1, and Q3 and 1.5 interquartile range (b) Comparison of the proportion of modified sites predicted as modified by m6Anet and by m6ACE across all 5-mers.

**Supplementary Figure 2.**
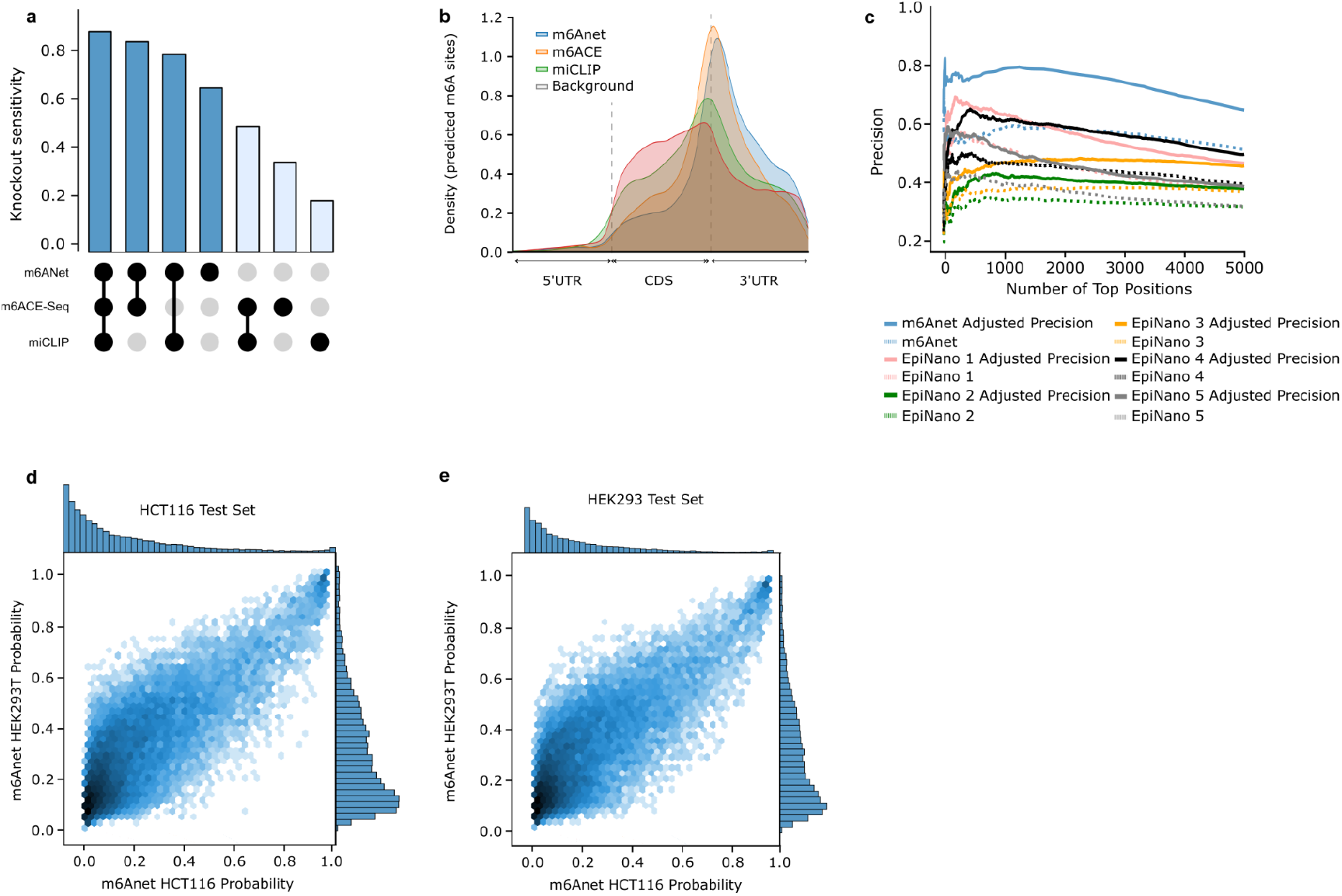
Performance comparison between *m6Anet*, m6ACE-seq and miCLIP on HEK293T cell line. (a) Percentage of captured sites that show significant shift in signal distribution against METTL3-KO for each of the three protocols (b) Metagene plot of the modified sites captured exactly by one of the three protocols against the background distribution of all DRACH sites in the data that has at least 20 readsTotal number of modified sites captured by *m6Anet*, m6ACE-seq and miCLIP (c) The adjusted true positive rate after including position sensitive to METTL3-KO of *m6Anet* and all 5 EpiNano models (d-f) Scatter plot of the predicted probability of the HEK293T model against the predicted probability of the HCT116 model on the HCT116 test set and HEK293T test set. The plot shows strong linear relationship between the prediction of the two models, indicating that *m6Anet* shows robustness in its prediction despite being trained on different cell lines.

**Supplementary Figure 3.**
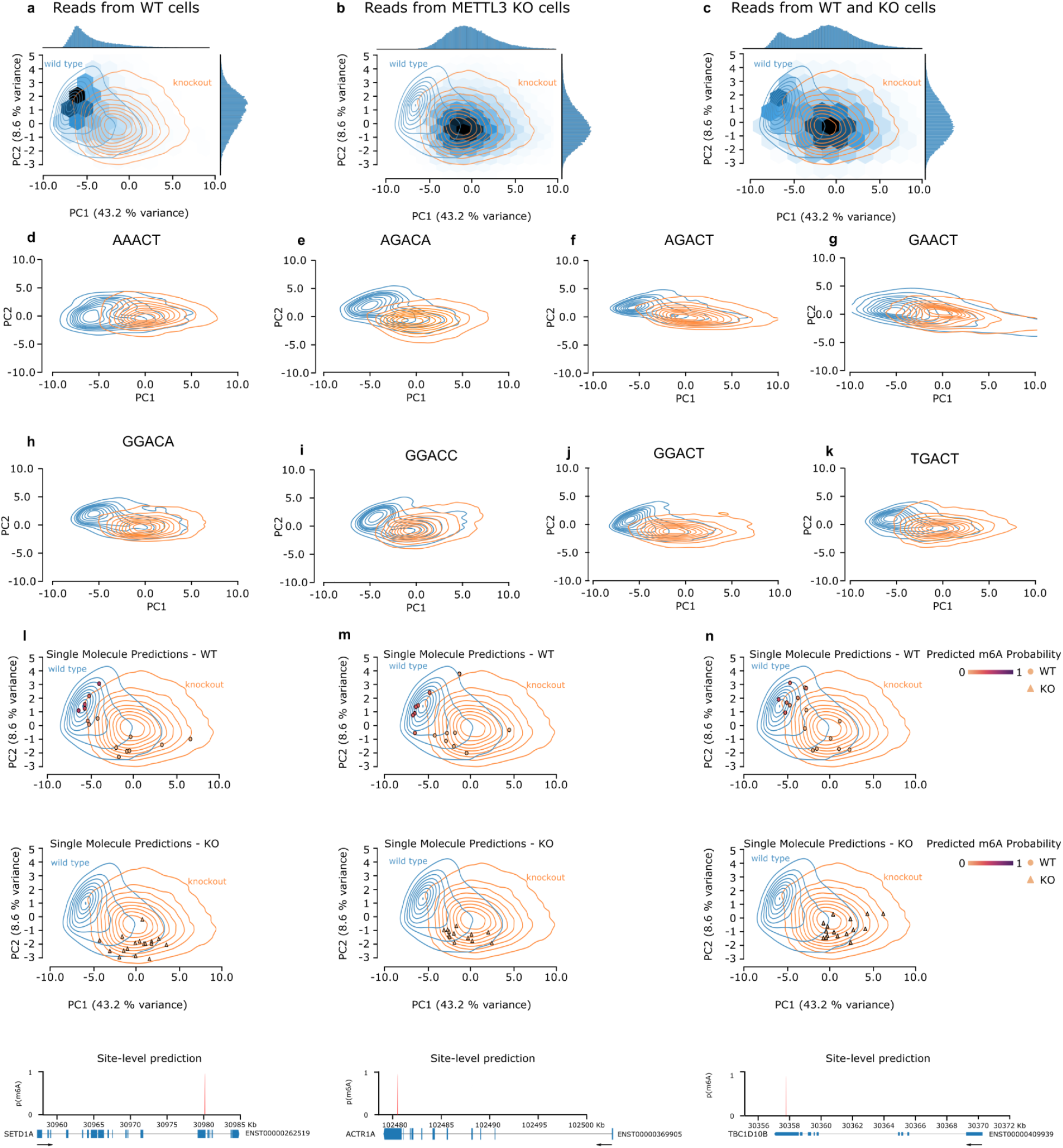
Quantification of m6Anet on HEK293T cell line. Experimental design: We sample 100 reads from each DRACH sites and extract the output from the second last layer of *m6Anet* for each of these reads and visualize them on two dimensional space using PCA (a) Hex plot of the top 4 modified 5-mer (GGACT, GAACT, GGACA, AGACT) for Wild Type sample, (b) Knockout sample (c) Both Wild-Type and Knockout sample after filtering for positions that are almost 100% modified on the Wild Type sample (p >= 0.9) and almost 0% modified on the KO sample (p <= 0.2) (d-f) Scatter plot of 20 randomly sampled reads from the second, third, and fourth ranked positions sorted by predicted modification probability on the Wild Type sample after the filter (g-n) Density plots of selected DRACH 5-mers that contain at least 20 modified sites (p >= 0.9 on WT samples) and at least 20 unmodified sites (p <= 0.2 on KO samples)

**Supplementary Figure 4.**
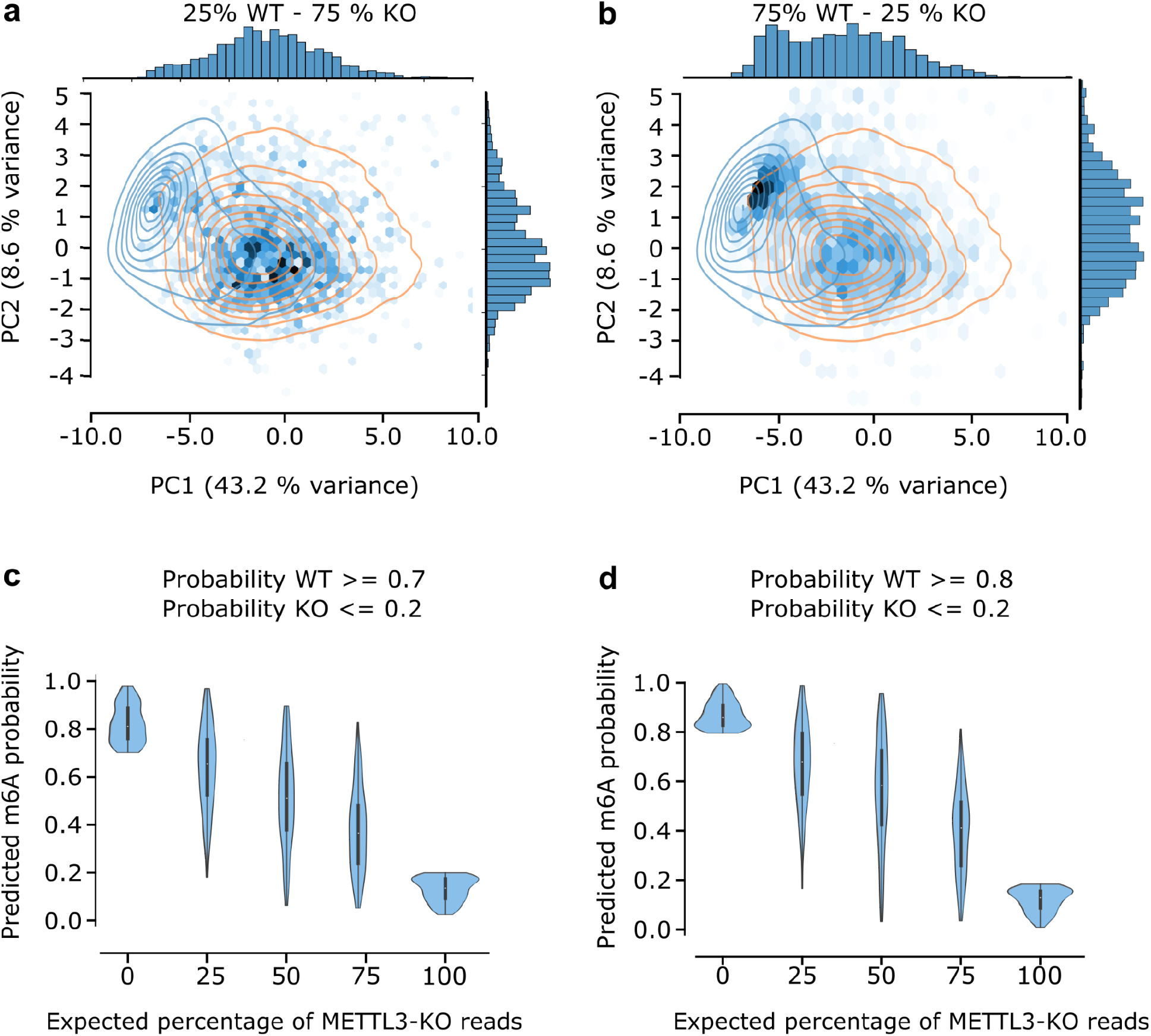
Changes in expected representation and predicted probability on the HEK293T Knockout Mixtures. Experimental design: We extract 100 reads from positions that show both high probability of modification (p >= 0.9) on the Wild Type sample and low probability of modification (p <= 0.2) on the corresponding KO sample and expressed across all 5 Wild Type - KO mixtures (a-e) Changes in the concentration of points as visualized on the first two principal components of the same PCA space as in Figure 4. The gradual shifts from (a) to (b) suggests that m6Anet read features capture the expected change in the stoichiometry of m6A modifications (c-d) Violin plot of the probability score of the top predicted positions by *m6Anet* across the 5 mixtures with less stringent requirement on the minimum probability of modification on the Wild Type samples. The plots still show the expected decrease in the predicted m6A probability as the percentage of METTL3-KO reads increase even with less stringent thresholds.

## Notes

https://github.com/GoekeLab/m6anet

### Competing Interest Statement

The authors have declared no competing interest.

